# Genome-Wide Association Testing for Haemorrhagic Bowel Syndrome in a Swiss Large White Pig Population

**DOI:** 10.1101/2024.04.05.588256

**Authors:** Arnav Mehrotra, Alexander S. Leonard, Cord Drögemüller, Alexander Grahofer, Negar Khayatzadeh, Andreas Hofer, Stefan Neuenschwander, Hubert Pausch

**Affiliations:** Animal Genomics, ETH Zürich, Universitätstrasse 2, Zürich, 8092, Switzerland; Institute of Genetics, Vetsuisse Faculty, University of Bern, Bern 3012, Switzerland; Clinic for Swine, Department for Clinical Veterinary Medicine, Vetsuisse Faculty, University of Bern, Bremgartenstrasse 109a, 3012, Bern, Switzerland; SUISAG, Allmend 10, 6204, Sempach, Switzerland; Animal Genetics, ETH Zürich, Tannenstrasse 1, 8092, Zürich, Switzerland

## Abstract

**Background:** The porcine haemorrhagic bowel syndrome (HBS) is a multifactorial disease causing fatal gastrointestinal disturbances and sudden death in fattening pigs. HBS is the leading cause of deaths during fattening in Swiss pigs, with unclear etiology. Environmental and management factors are associated with HBS incidence, but recent findings also suggest a potential genetic predisposition. Pigs sired by a Swiss Large White (SLW) line appear more prone to HBS. Here we conduct genome-wide association studies (GWAS) for HBS between cases and controls to investigate potential genetic factors for the disease in Swiss fattening pigs.

**Results:** Our study included 1,036 HBS cases and 4,080 controls with available microarray genotypes or whole-genome sequencing data. Variant positions were determined according to the current porcine reference assembly (Sscrofa11.1) or a HiFi-based SLW haplotype assembly which we constructed using trio-binning. GWAS for HBS were conducted using 12.49 to 15.46 million biallelic variants in three mapping cohorts consisting of purebred animals from SLW sire and dam lines, or crosses between these two parental lines. The statistical model applied for the GWAS accounted for animal relatedness, population structure, and an imbalanced case/control ratio. No sequence variants significantly associated with HBS were identified, regardless of the cohort analysed and the reference sequence considered.

**Conclusions:** The lack of genetic associations despite a relatively large sample size suggests that susceptibility to HBS in the studied SLW population is not due to large effect variants but may be influenced by numerous small effect genetic variants, in addition to environmental and management factors.

## Background

Haemorrhagic Bowel Syndrome (HBS) in pigs is a multifactorial disease that has become a substantial threat to swine production. The disease manifests in fatal gastrointestinal disturbances and sudden death in finisher pigs [1]. Key characteristics of HBS include pallor and abdominal distention in carcasses, with intestinal torsion frequently observed during necropsy [2,3]. Despite its substantial impact on animal welfare and the economics of pig production, the etiology of HBS is not fully understood. Several factors contributing to the incidence of the disease have been identified, including feeding systems [4], the origins of the fattening pigs, cleaning frequency of feed distribution systems, and the width of feeding places [2,5]. Seasonal variations [1,3,6], antimicrobial usage in feed [2,7], gut microflora composition [3], and infectious agents such as *Clostridium perfringens* and *Enterobacteriaceae* [2,4] have also been implicated.

A genetic component to HBS susceptibility has been proposed by previous studies [1,4]. A recent study by Holenweger et al. [5] also revealed that fattening pigs descending from a Swiss Large White (SLW) sire line were overrepresented among affected animals. These recent findings suggest that genetic variants may contribute to the susceptibility to HBS and may explain at least part of the observed across-breed variability to develop the disease.

This study aimed at identifying genetic variants associated with HBS in Swiss pigs through genome-wide association testing, thereby contributing to a better understanding of the etiology of the syndrome and facilitating the development of breeding strategies to prevent disease incidence.

## Methods

Tissue samples of 1036 pigs that died from HBS were collected on farm and at animal carcass collection points in Switzerland over a period of six months, through SUISAG, the service partner for Swiss pig producers. HBS was confirmed post-mortem based on the inspection of the intestinal tract by stock veterinarians and veterinarians from SUISAG-SGD. We prepared DNA from ear tissue samples of 1036 HBS-affected pigs using the Promega Maxwell RSC DNA System (Promega, Dübendorf, Switzerland) and sent it to Gencove for low-pass sequencing (https://gencove.com/). An average number of 945K (between 150K and 5.15M) read pairs (2 × 150bp) corresponding to an average genome coverage of approximately 1-fold were collected for the HBS cases. Genotypes for 45,100,556 biallelic sequence variants (SNP) corresponding to the Sscrofa11.1 (GCA_000003025.6) reference sequence [8] were provided by Gencove. To infer the population structure, we extracted a subset of 48,919 SNPs from the genotypes that are also present on the SNP arrays routinely used for genomic prediction in Switzerland. These SNP genotypes were combined with array-derived genotype data of 17,006 pigs that were genotyped for routine genomic prediction, resulting in a combined dataset of 18,042 animals. After removing 4,554 markers with minor allele frequency (MAF) less than 5% using PLINK (v1.90) [9], a genomic relationship matrix (GRM) was constructed based on 44k SNPs using GCTA (v1.94.1) [10] and the top principal components were extracted and visualized to assess the structure of the genotyped populations and select three ancestry-matched control cohorts. The cohorts consisted of 1) purebred animals from a SLW sire line, (70 cases, 280 controls), 2) purebred animals from a SLW dam line (61 cases, 244 controls), and 3) a mixed population of animals from the sire and dam lines, their crosses, as well as crosses between either the sire or the dam line and Landrace animals (1,020 cases, 4,080 controls), maintaining a 1:4 case-to-control ratio in all cohorts. Boars and sows genotyped for routine genomic prediction whose progeny were registered in the national herdbook and therefore not affected by HBS before reaching reproductive age were considered as control animals.

Genotypes for the control animals were imputed to the whole-genome sequence level with Beagle v5.4 [11] using a sequenced reference panel of 421 pigs that included mostly SLW animals. The reference animals had genotypes at 22,018,148 autosomal and 350,478 X-chromosomal sequence variants [12]. We retained 15.43 million autosomal 192,930 X-chromosomal biallelic SNPs that had a model-based accuracy of imputation greater than 0.8. Sex of HBS cases was inferred based on allele frequency estimates of X-chromosomal markers (--impute-sex flag in PLINK).

We considered SNPs with a minor allele frequency greater than 5% and less than 10% missingness for the subsequent association tests. For the purebred cohorts we excluded SNPs from genome-wide association testing that deviated significantly (p<1 × 10^−6^) from Hardy-Weinberg proportions. After the quality control, we retained between 12.49 and 13.46 million SNP for association testing in the three cohorts.

The genome-wide association studies (GWAS) in the three cohorts were performed using a generalized linear mixed model implemented in SAIGE (v1.0.9) [13] which accounts for sample relatedness and an imbalanced ratio of cases and controls. The top 10 principal components derived from a GRM, and sex were included as fixed factors, and a GRM was fitted as random effect. A Bonferroni-corrected significance threshold (p=3.1 × 10^−9^) was applied to account for multiple testing.

A haplotype of the SLW breed was assembled through trio-binning [14], utilizing PacBio high-fidelity (HiFi) reads from a male offspring originating from a crossing between a purebred Swiss Large White boar and an Alpenschwein sow. High Molecular Weight (HMW) DNA was extracted from a liver tissue sample of the F1 using the Monarch® HMW DNA Extraction kit and sequenced on the PacBio Sequel IIe platform, employing three SMRT cells to generate 4.93 million HiFi reads with an average length of 17.73 kb. Additionally, DNA from maternal blood and paternal liver tissue samples was sequenced on an Illumina Novaseq6000 to generate short reads with an average coverage of 28.7x. Haplotype-resolved assemblies were then generated with hifiasm v0.16.1 [15] as outlined in [16]. The paternal haplotype (SLW) assembly size was 2.406 Gb across 998 contigs, with an average Quality Value (QV) of 49.96 and a contig N50 of 30.62 Mb. The assembly achieved a BUSCO single-copy score of 87.9%.

Paired-end short reads from our imputation reference cohort consisting of 421 samples were aligned to the herein generated SLW assembly using bwa-mem2 [17]. Variant calling was then performed with DeepVariant v1.4 and GLnexus v1.4.1 [18] resulting in 29.73 million autosomal variants. From these, an imputation reference panel was constructed using 24.75 million biallelic SNPs using the bcftools v1.6 [19] view command with the ‘–m2 – M2 –v snps’ flags. We imputed the low-pass sequencing data of HBS cases to the sequence level with Glimpse v1.1 [20]. Coordinates of SNP array genotypes of controls were lifted over to the SLW assembly using nf-LO [21] and subsequently imputed up to the sequence level with Beagle v5.4 [11]. The imputed case and control datasets were merged, retaining 19.78 million SNPs with a missingness of less than 10% and a MAF greater than 0.05 for GRM preparation and principal component analysis. Following quality control performed separately for each cohort, as previously detailed (--maf 0.05, --geno 0.1; --hwe 1 × 10^-6 for purebred cohorts), we retained between 13.71 and 15.46 million SNPs for GWAS with SAIGE, as described earlier.

## Results and discussion

Genome-wide association studies with partially imputed sequence variant genotypes were performed to identify genomic loci associated with susceptibility to HBS in Swiss fattening pigs. A principal components analysis identified three distinct cohorts for association testing: purebred animals from the SLW sire line, purebred animals from the SLW dam line, and crosses (Figure 1A, 1D, 1G). The majority of HBS cases fell into the latter cohort, as it included the fattening pigs which were produced by the crossing of purebred parental lines.

**Figure 1.**
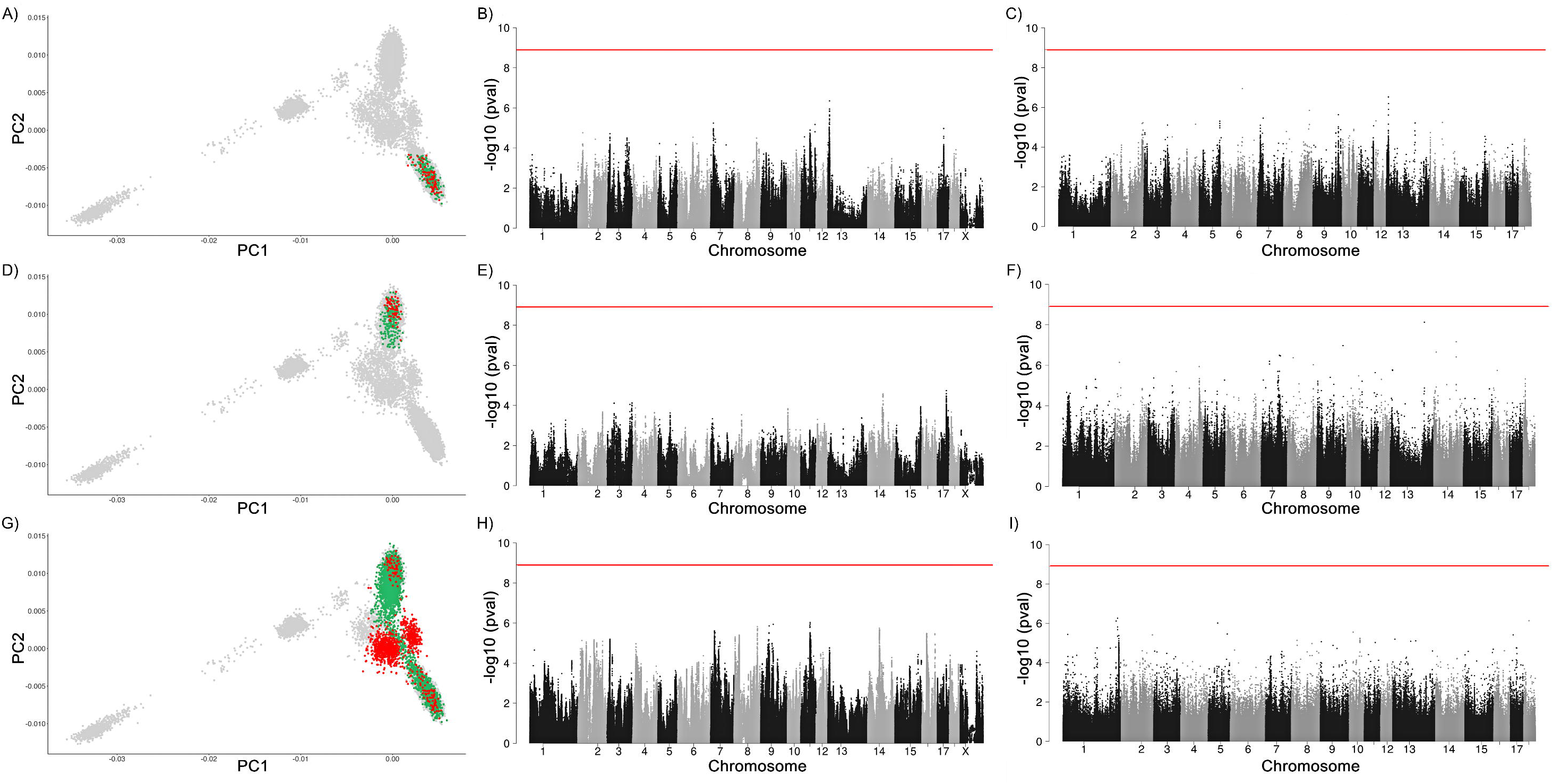
Principal component analysis (PCA) and genome-wide association study (GWAS) results across three cohorts. PCA plots highlighting cases (red) and controls (green) selected for the three cohorts: purebred Swiss Large White sire line (A), purebred Swiss Large White dam line (D), and a mixed population including these two lines their crosses, and crosses between either of those lines and Landrace (G). The corresponding Manhattan plots (B, E, H) illustrate the GWAS results using the Sscrofa11.1 reference genome. Plots (C, F, I) depict GWAS results utilizing the herein established Swiss Large White reference assembly. The red line across all Manhattan plots represents the Bonferroni-corrected significance threshold.

Our initial case/control association analyses relied on between 12.49 and 13.46 million variants for which genotypes were called from alignments against the current Sscrofa 11.1 genome assembly which was assembled from a Duroc boar [8]. Neither the within-breed nor the multi-breed GWAS revealed markers significantly associated with HBS (Figure 1B, 1E, 1H). The choice of the reference assembly can impact sequence read alignment, variant calling, and downstream genetic analyses [22], particularly if the target breed diverged from the reference sequence. We repeated the GWAS with between 13.71 and 15.46 million SNPs for which genotypes were called from a SLW assembly (Fig 1C, 1F, 1I), but again, found no significant genetic variants associated with HBS in either of the cohorts tested. The lack of significant associations is somewhat surprising as previous studies identified the SLW sire line as the main risk factor for developing HBS [5]. Given a relatively low effective population size of 72 and 44 for the sire and dam lines [23], respectively, and the fact that both populations diverged less than 20 years ago, we are confident that our cohort was large enough to identify large effect genetic variants for developing HBS. Therefore, our findings suggest that the previously reported breed-specific predisposition to HBS is mainly driven by small effect genetic variants, which are possibly modulated by environmental and management factors.

Incorporating environmental variables into the case/control GWAS was not possible as the case and control groups were reared under disparate conditions. Future research efforts to better understand the genetic architecture for the development of HBS in fattening pigs and to identify associated genetic variants should ideally select control pigs from the same fattening farms as the cases, ensuring that these controls are closely matched to the cases in terms of age and weight profile and have successfully reached slaughter. Unfortunately, implementing such an effort is not straightforward as fattening pigs are not routinely genotyped. Farms with high and low incidences of HBS identified before [5] could serve as candidates for undertaking such genotyping efforts. However, considering that an average Swiss fattening farm produces approximately 500 fattening pigs per year, and assuming an HBS incidence of 1% [5], the sampling of a case/control cohort with sufficient statistical power to identify trait-associated small effect variants appears to be a major undertaking. Nevertheless, such a cohort may also serve as a reference for the implementation of genomic prediction.

The lack of pedigree information for the mostly cross-bred HBS cases, together with the structured and related case/control cohorts, makes it difficult to estimate heritability of HBS. Although the statistical methods implemented in SAIGE and fastGWA-GLMM [24] minimize type I errors attributable to relatedness and imbalances between cases and controls in GWAS, the variance components they estimate through penalized quasi-likelihood are not precise enough for calculating heritability [13, 24]. Documenting pedigree information for fattening pigs could contribute to a better understanding of the genetic architecture underlying the disease.

Finally, the potential impact of structural variations (SVs), which may not be adequately tagged by the SNPs used in our study, should also be considered. Although the newly built, breed-specific SLW assembly makes variants overlapping insertions with respect to the Duroc-based reference sequence amenable to association mapping, the short-read sequencing-based approach doesn’t allow to comprehensively study SVs [25]. The establishment of a porcine SV imputation reference panel for pigs could enable future HBS research by integrating SV data for a more comprehensive association analysis.

## Conclusions

We report the first GWAS for HBS in Swiss pigs reported to be at risk for HBS, a sporadically occurring disorder characterized by sudden death in fattening pigs. Our comprehensive genetic analysis, spanning several breeds and two different reference genomes, did not reveal any significant genetic markers associated with HBS. This finding suggests that the genetic susceptibility to HBS is likely to involve small effect genetic variants and/or more complex SVs that may interact with environmental and management factors, rather than large effect genetic variants. Future research should prioritize the selection of cases and controls from the same fattening farms, allowing a clearer distinction between genetic predisposition and environmental influences.

## List of abbreviations

BUSCO: Benchmarking Universal Single-Copy Orthologs
GRM: Genomic Relationship Matrix
GWAS: Genome-Wide Association Studies
HBS: Haemorrhagic Bowel Syndrome
HiFi: High Fidelity
HMW: High Molecular Weight
MAF: Minor Allele Frequency
QV: Quality Value
SLW: Swiss Large White
SNP: Single Nucleotide Polymorphism
SV: Structural Variation

## Declarations

### Ethics approval and consent to participate

Tissue samples for the HIS cohort were collected from dead pigs. The sampling of blood from the F1 and its parents used for constructing the de novo assembly was approved by the veterinary office of the Canton of Zurich (animal experimentation permit ZH077/2022).

### Consent for publication

Not applicable

### Availability of data and materials

Low-pass sequencing data of pigs affected by HBS have been deposited at the European Nucleotide Archive (ENA) of the EMBL at BioProject PRJEB62539. Raw sequence read data of pigs used to prepare the imputation reference panel are available at the European Nucleotide Archive (ENA) of the EMBL at BioProjects PRJEB38156 and PRJEB39374. Long and short sequencing reads from the trio used to generate the SLW assembly are at PRJEB74562.

### Authors’ contributions

AM performed the analyses and wrote the first draft of the manuscript with input from ASL and HP. ASL assembled the SLW haplotype. SN collected samples from the trio used for SLW assembly. HP and AH conceptualized the study. CD, AG, AH, NK and HP conceptualized the project. All authors read and approved the final manuscript.

## Acknowledgements

We acknowledge support from Hans-Peter Grünenfelder, Tanja Kutzer, Thomas Echtermann and Xaver Sidler to produce the F1 cross. We thank the Functional Genomics Center Zurich (Anna Bratus-Neuenschwander) for generating HiFi data. The assistance of Nathalie Besuchet-Schmutz with DNA extraction is acknowledged.

